## Supplementary material for "The transcriptional and epigenetic reprogramming associated with chronic IL1β-mediated expansion and myeloid priming in TET2 deficient stem and progenitor cells": Figures

Supplementary Figure 1

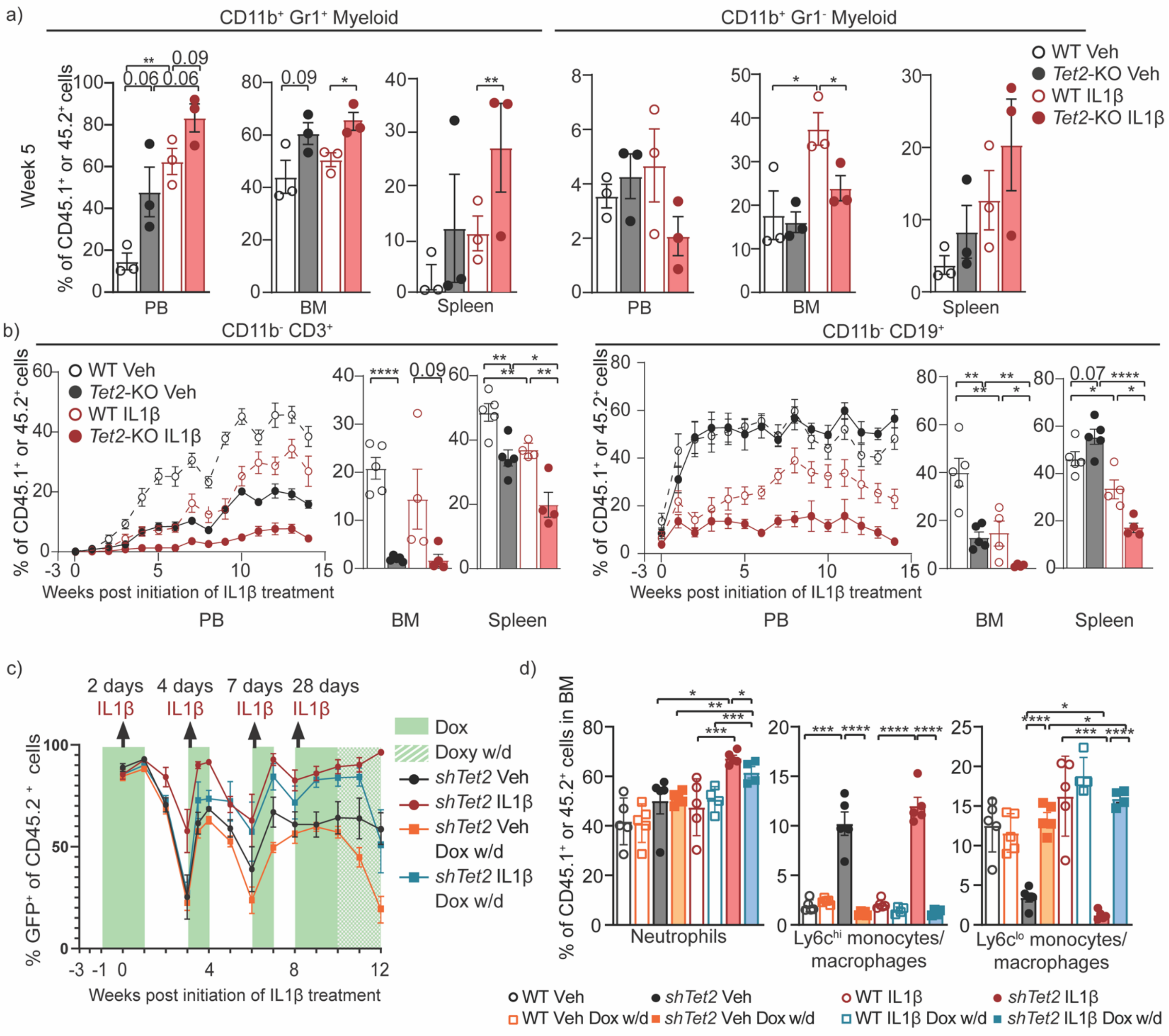


**Supplementary Figure 1: Chronic IL1β exposure enhances myelopoiesis at the expense of lymphoid cell frequency.**

**(a-b)** Lineage-depleted BM cells from WT CD45.1 and *Tet2*-KO CD45.2 mice were transplanted into lethally irradiated WT CD45.1/2 mice and after 3 weeks treated with IL1β (500 ng/mouse/day) or vehicle daily. **(a)** The frequency of myeloid subsets (CD11b$^+$Gr1$^{hi}$, CD11b$^+$Gr1$^{lo}$) at week 5 (week 15 is shown in main Fig. 1, n = 3 mice/group) and **(b)** T cells (CD11b$^-$ CD3$^+$) and B cells (CD11b$^-$CD19$^+$) at 15 weeks in PB, BM, and spleen (n = 4-5 mice/group). **(c-d)** Lineage-depleted BM cells from wild-type (WT) CD45.1+ and Rosa-rtTA-driven inducible *Tet2* knockdown (*shTet2*) mice were transplanted into lethally irradiated WT CD45.1/2 mice, which were treated with doxycycline (dox) two weeks after transplantation and IL1β or vehicle three weeks after transplantation for increasing intervals up to 10 weeks and then with and without doxycycline (withdrawal, w/d) for an additional two weeks (Fig. 1d), analyzed by flow cytometry (n = 4-5 mice/group) (described in Fig. 1d). **(c)** percentage of CD45$^+$ *shTet2* which are GFP$^+$ in PB **(d)** and the percentage of neutrophils (CD45$^+$CD11b$^+$Ly6c$^{int}$Ly6g$^+$), ly6c$^{hi}$ monocytes/macrophages (CD45$^+$CD11b$^+$Ly6c$^+$Ly6g$^-$) and ly6c$^{lo}$ monocytes/macrophages (CD45$^+$CD11b$^+$Ly6c$^-$Ly6g$^-$) in the BM (n = 4-5 mice/group). Error bars represent mean ± SEM. Student's two-tailed t-Test was used to determine significance: *p < 0.05, **p < 0.01, ***p < 0.001, ****p < 0.0001.

Supplementary Figure 2

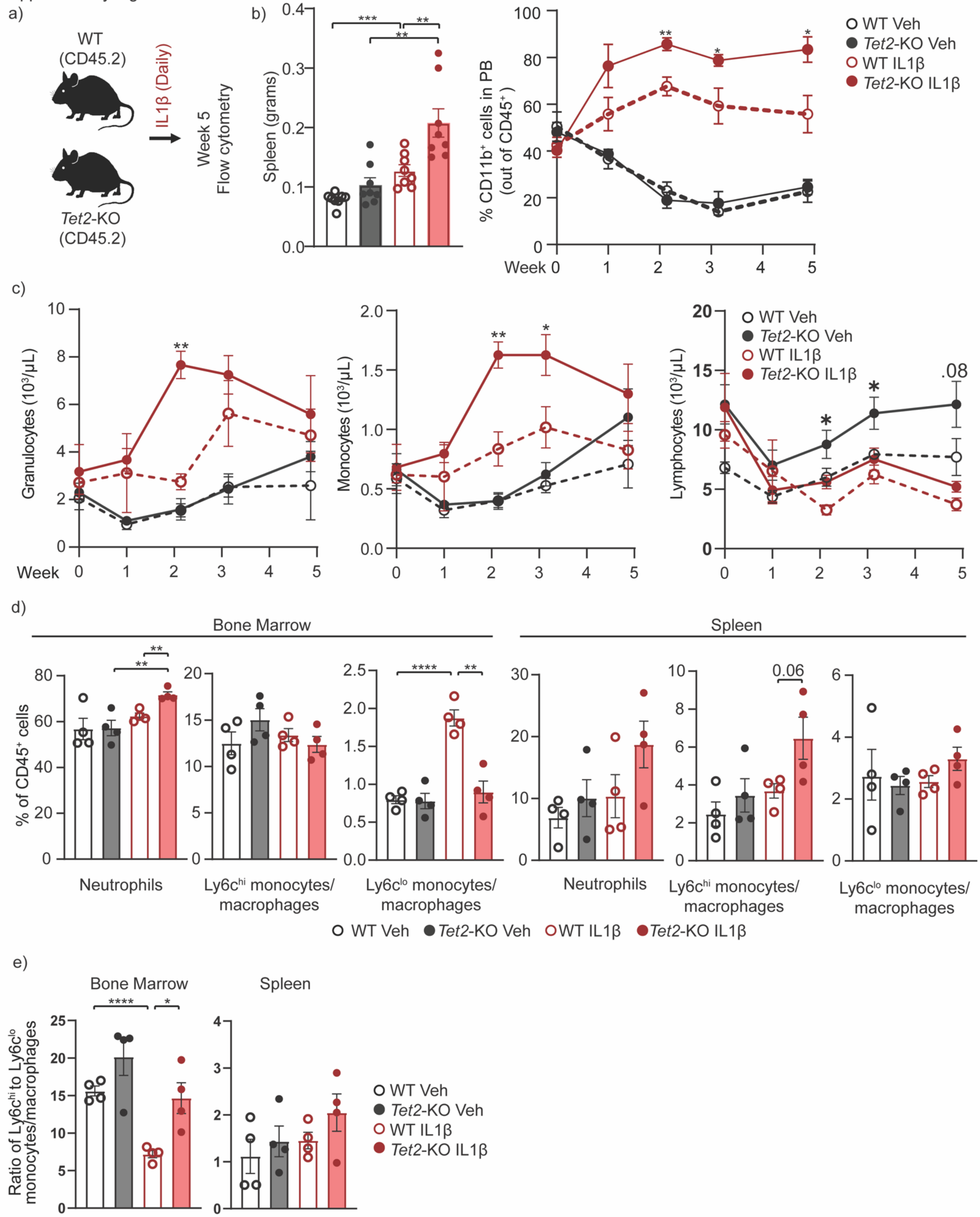

**Supplementary Figure 2: Chronic IL1β exposure enhances myeloid expansion of *Tet2*-KO cells in non-competitive murine models.**

*Tet2*$^{fl/fl}$ (WT) and *Vav-Cre Tet2*$^{fl/fl}$ (*Tet2*-KO) mice were treated with IL1β (500 ng/mouse/day) or vehicle ($n = 4$ mice/group) for five weeks and peripheral blood (PB), bone marrow (BM) and spleen samples were analyzed by flow cytometry. **(a)** Experimental design, **(b)** spleen weight in grams (left) and the percentage of the PB CD45$^{+}$ cells which are myeloid (CD45$^{+}$CD11b$^{+}$, right). **(c)** Differential blood cell counts representing granulocytes, monocytes, lymphocytes, hematocrit, hemoglobin, and platelets in the PB. **(d)** Neutrophils, ly6c$^{hi}$ monocytes/macrophages and ly6c$^{lo}$ monocytes/macrophages and **(e)** ratio of Ly6c$^{hi}$ to ly6c$^{lo}$ monocytes/macrophages in BM and spleen cells at week five by flow cytometry. Error bars represent mean ± SEM. Student's two-tailed t-Test was used to determine significance: *$p < 0.05$, **$p < 0.01$, ***$p < 0.001$, ****$p < 0.0001$.

Supplementary Figure 3

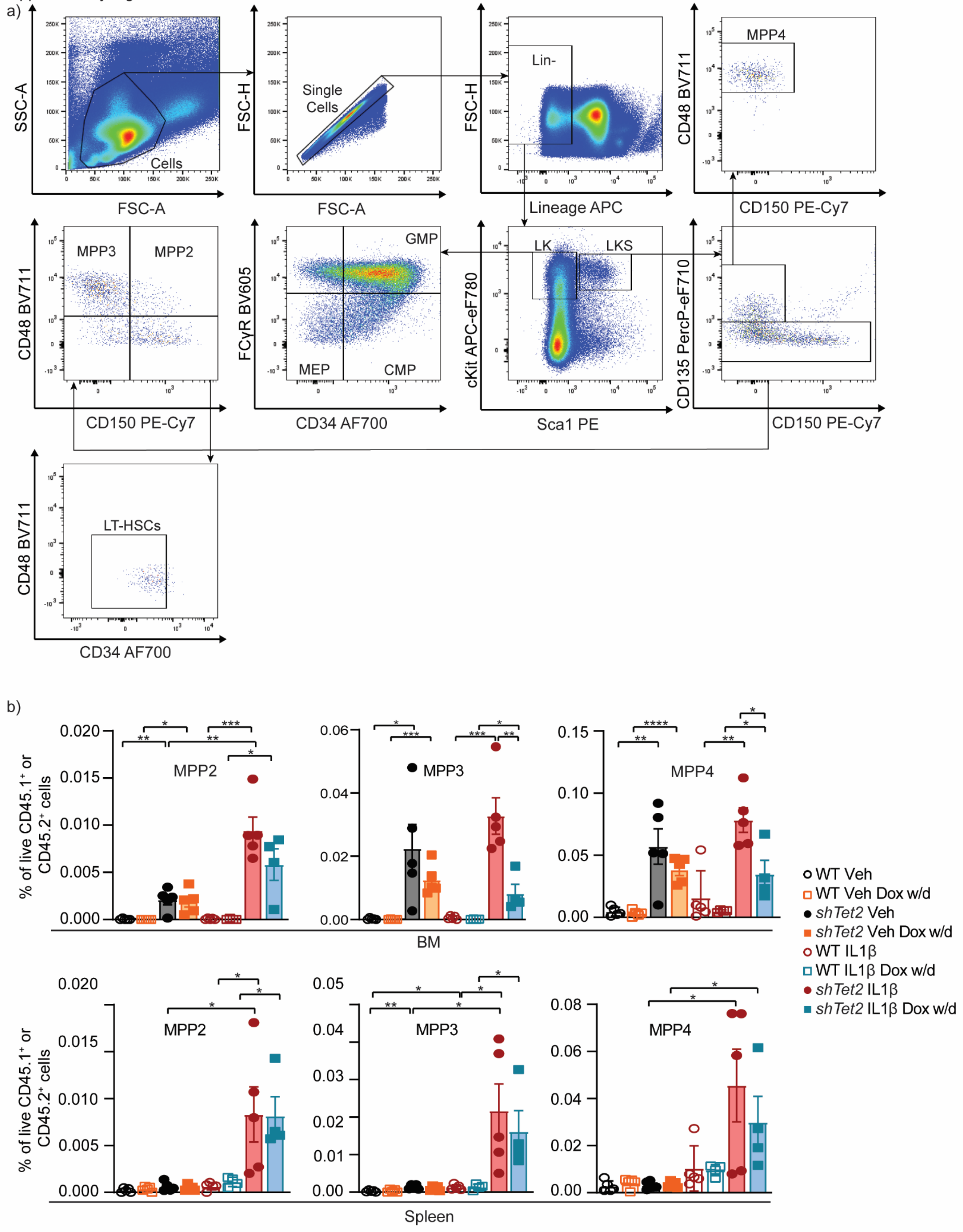

**Supplementary Figure 3: IL1β stimulation promotes HSPC expansion in mice with *Tet2* knockdown relative to WT in a competition repopulation experiment.**

**(a)** Representative gating scheme for HSPCs for flow cytometric analysis of the BM from a WT mouse. **(b)** Lineage-depleted BM cells derived from WT CD45.1 and *shTet2* CD45.2 mice transplanted into lethally irradiated WT CD45.1$^{+}$CD45.2$^{+}$ mice, which were treated with IL1β or vehicle as well as with and without doxycycline (withdrawal; dox w/d) as described in Fig 1d and analyzed by flow cytometry at the endpoint. Percentage out of live cells gated out of WT CD45.1 and *shTet2* CD45.2 for MPP2s, MPP3s, and MPP4s within spleen and BM of recipient mice (n = 4-5 mice/group). Error bars represent mean ± SEM. Student's two-tailed t-Test was used to determine significance: *$p < 0.05$, **$p < 0.01$, ***$p < 0.001$, ****$p < 0.0001$.

Supplementary Figure 4

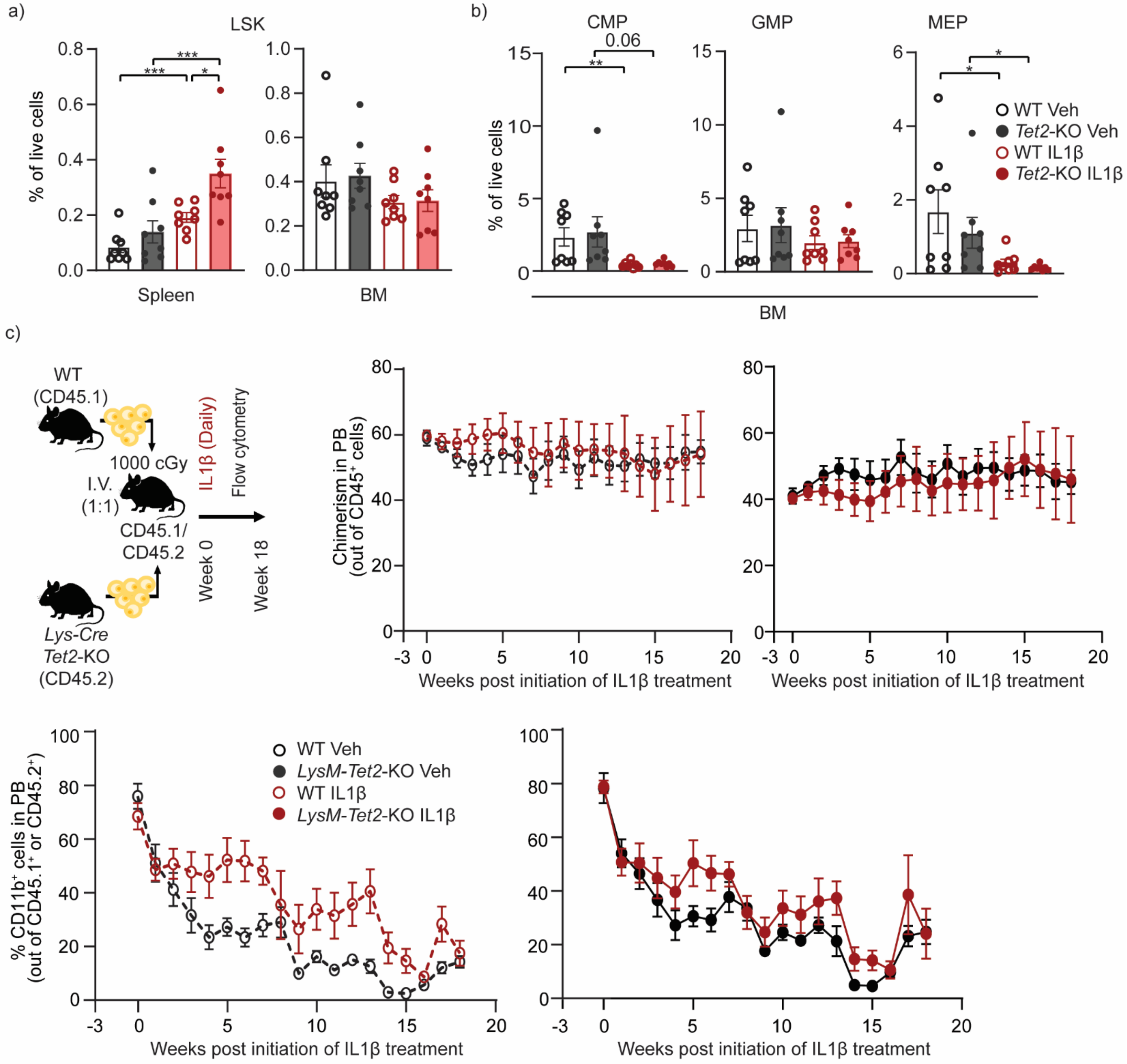


**Supplementary Figure 4: IL1β stimulation promotes myeloid bias of *Tet2*-KO CMPs and deletion of *Tet2* within HSPCs is necessary for their expansion and myeloid bias.**

**(a)** WT and *Tet2*-KO mice were treated with IL1β or vehicle for five weeks (Supplementary Fig. 1a) and analyzed by flow cytometry for the percentage of live $CD45^+$ cells which are LSKs in the spleen or BM or **(b)** CMPs, GMPs, and MEPs in the BM. (**c**) Lineage-depleted BM cells were harvested from WT CD45.1 and *LysM-Cre Tet2*$^{-/-}$ CD45.2 (*LysM-Cre Tet2*-KO) mice into lethally irradiated WT $CD45.1^+CD45.2^+$ mice which after 3 weeks were treated with IL1β or vehicle. The percentage of WT CD45.1 and *LysM-Cre Tet2*-KO CD45.2 cells out of all $CD45^+$ cells (chimerism) and the percentage WT CD45.1 and *Tet2*-KO CD45.2 cells which are myeloid in the PB. Error bars represent mean ± SEM. Student's two-tailed t-Test was used to determine significance: *p < 0.05, **p < 0.01, ***p < 0.001.

Supplementary Figure 5

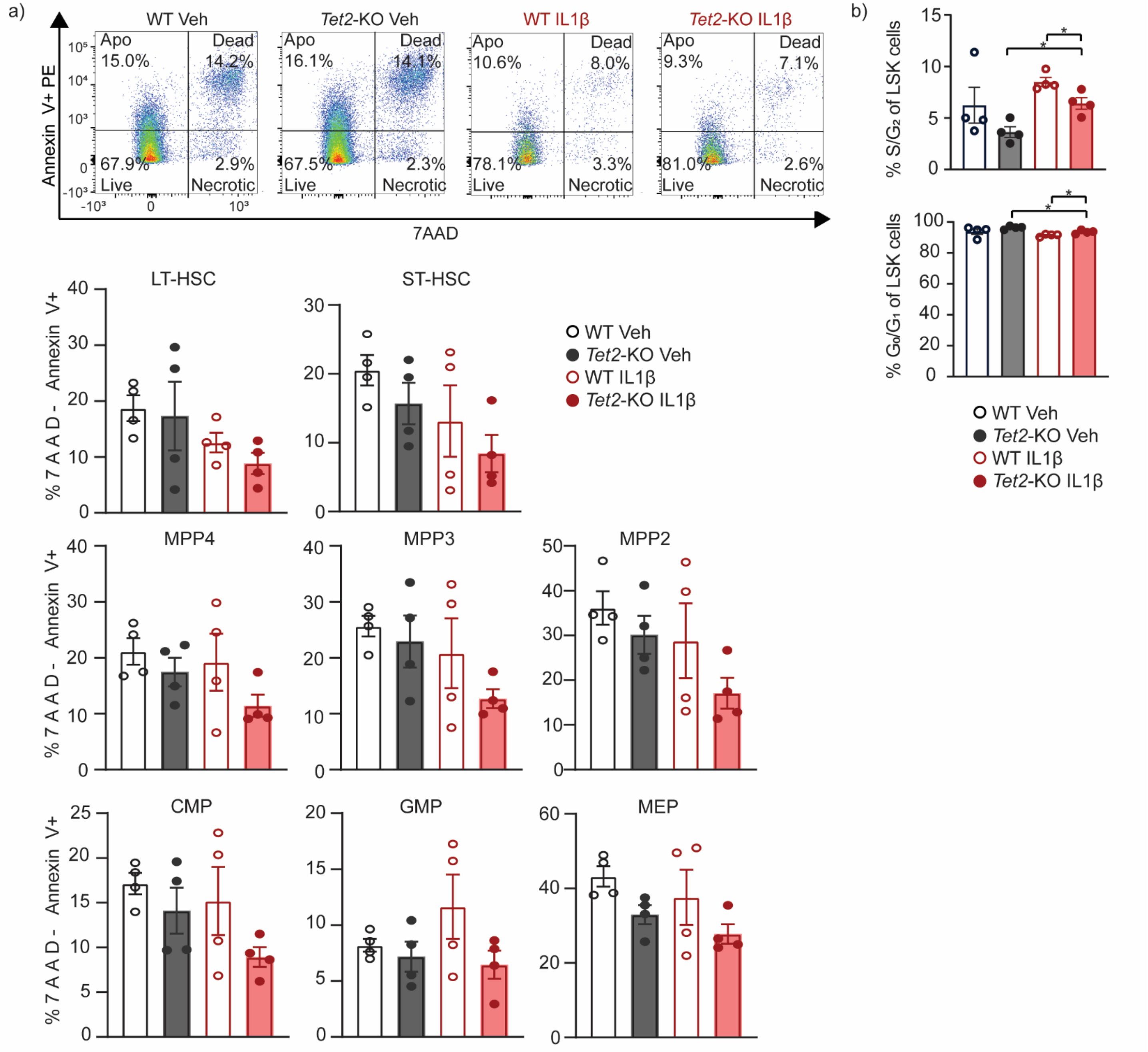


**Supplementary Figure 5: IL1β-mediated expansion of *Tet2*-KO HSPCs is associated with reduced S/$G_2$ frequency without significant differences in the frequency of apoptosis.**

**(a-b)** WT and *Tet2*-KO mice were treated with IL1β for 5 weeks (as described in Supplementary Fig. 2a; n = 4 mice/group). The percentage of apoptotic ($7AAD^{-}$Ann $V^{+}$) cells for HSPC subsets was quantified by flow cytometry. **(b)** Flow cytometry analysis of LSK cells isolated from the BM of *Tet2*-KO and WT mice treated with and without IL1β for 5 weeks, with cell cycle proportions using KI-67 and DAPI (right, n = 4 mice/group). Error bars represent mean ± SEM. Student's two-tailed t-Test was used to determine significance: *$p < 0.05$.

Supplementary Figure 6

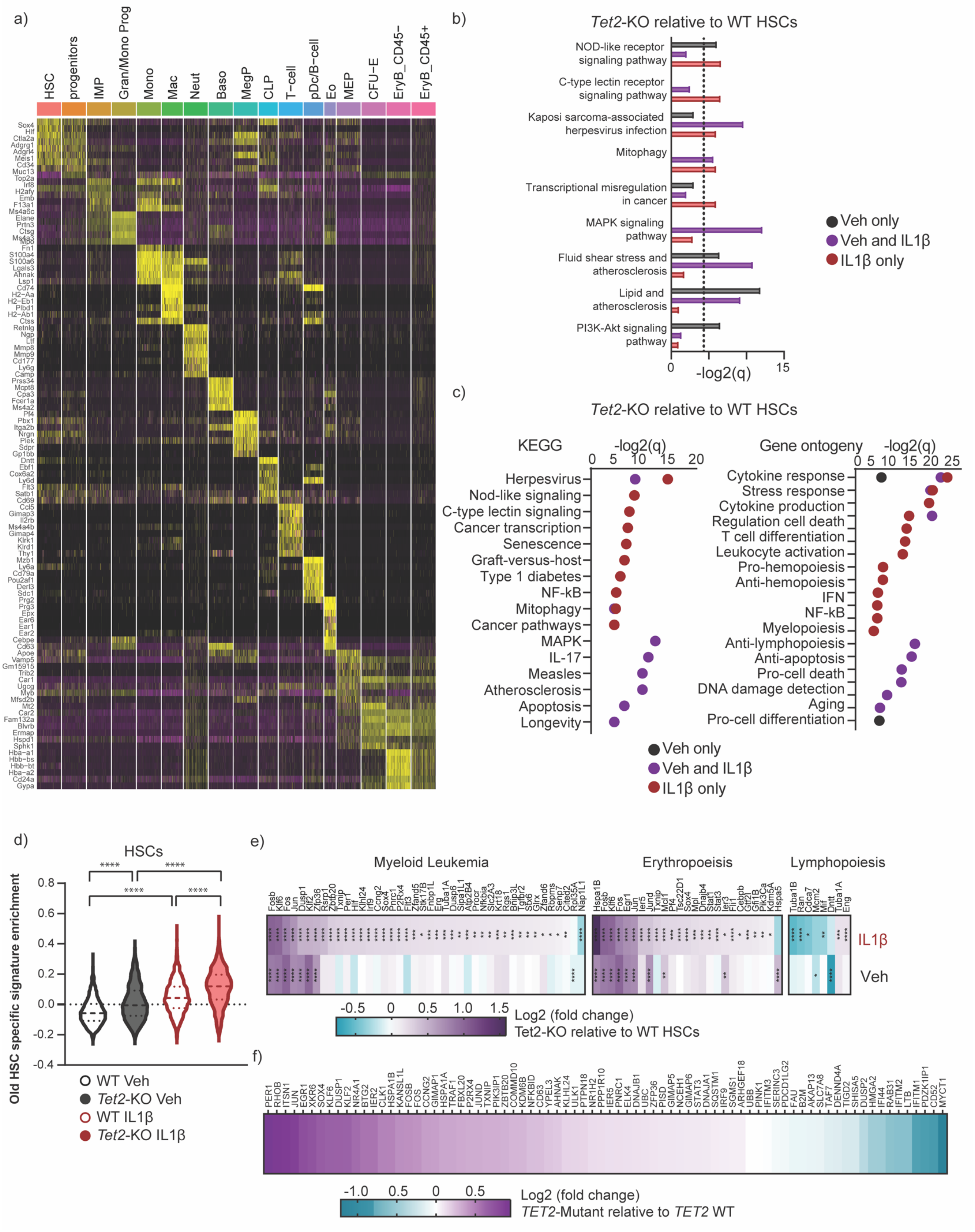

**Supplementary Figure 6: 10x single cell RNA sequencing to identify cellular clusters from *Tet2*-KO and WT mice with and without IL1β stimulation *in vivo*.**

**(a)** Heatmap showing gene expression of cluster defining-genes for individual cells from each cluster. (**b)** Enrichr ontology (KEGG 2021 human) of DEGs upregulated in *Tet2*-KO relative to WT HSCs in vehicle alone (black), IL1β alone (red) or in both (purple) with q values calculated by Enrichr. **(c)** STRING KEGG response to cytokine from biological process gene ontology (GO:0034097) and MAPK signaling pathway from KEGG Pathways (mmu04010) of DEGs upregulated in *Tet2*-KO relative to WT only under vehicle alone (black), IL1β alone (red), or both conditions (purple) with q values calculated by STRING. **(d)** Enrichment of transcriptional signature specific to old HSCs (Kirschner et al., Cell Rep, 2017) within HSCs. **(e)** Heatmap of upregulated DEGs in *Tet2*-KO relative to WT HSCs with and without IL1β stimulation. Genes shown are the lead genes for enriched pathways by GSEA analysis, with genes identified in multiple pathways shown multiple times. q values were determined by DESeq2. **(f)** Differential gene expression in *TET2*-mutant relative to *TET2*-WT de novo primary AML for the top 100 human orthologous genes that are upregulated in *Tet2*-KO relative to WT murine HSCs stimulated with IL1β. For q values: *$q < 0.05$, **$q < 0.01$, ***$q < 0.001$, ****$q < 0.0001$.

Supplementary Figure 7

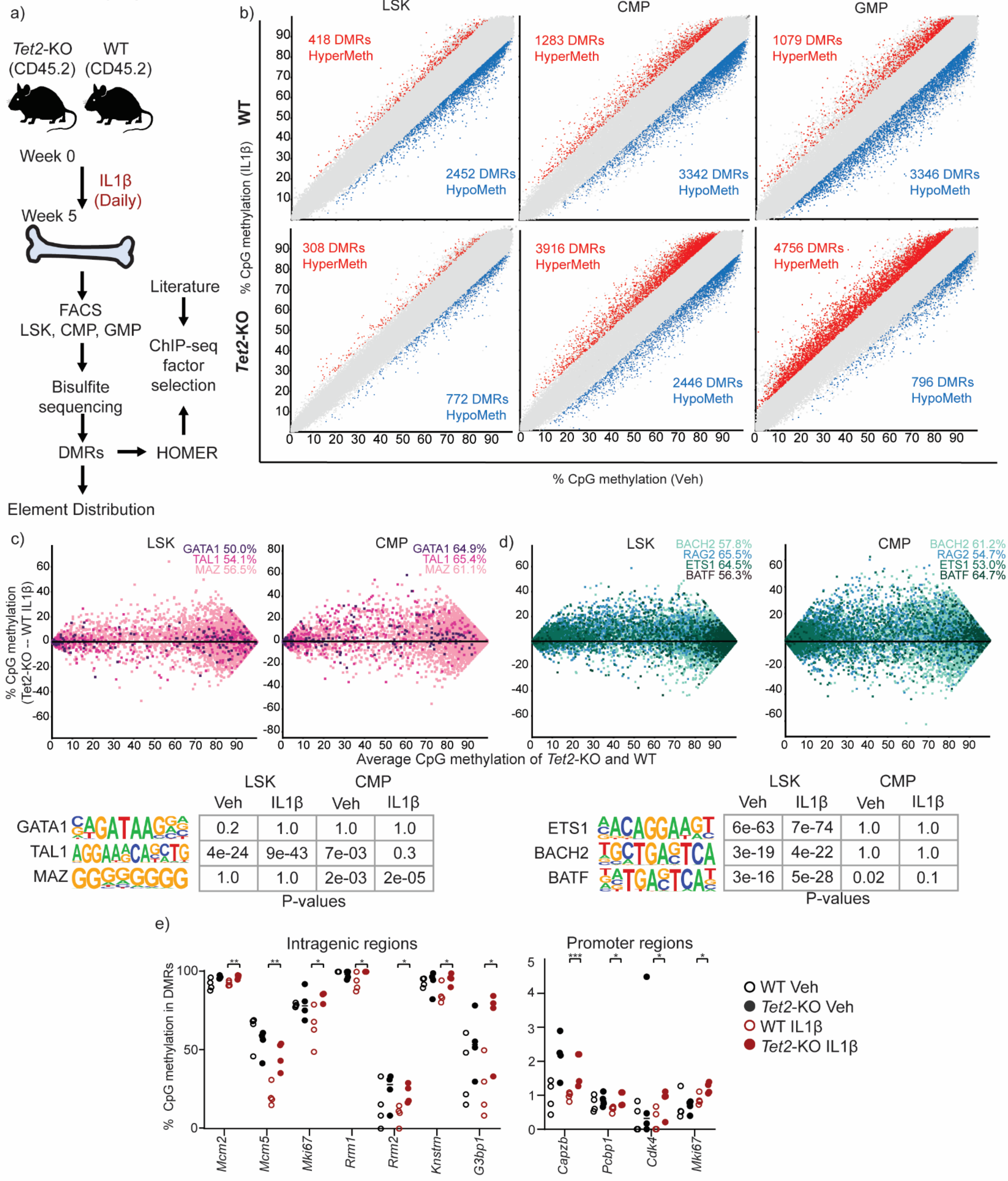


| | LSK Veh | LSK IL1β | CMP Veh | CMP IL1β |
|---|---|---|---|---|
| GATA1 | 0.2 | 1.0 | 1.0 | 1.0 |
| TAL1 | 4e-24 | 9e-43 | 7e-03 | 0.3 |
| MAZ | 1.0 | 1.0 | 2e-03 | 2e-05 |

P-values

| | LSK Veh | LSK IL1β | CMP Veh | CMP IL1β |
|---|---|---|---|---|
| ETS1 | 6e-63 | 7e-74 | 1.0 | 1.0 |
| BACH2 | 3e-19 | 4e-22 | 1.0 | 1.0 |
| BATF | 3e-16 | 5e-28 | 0.02 | 0.1 |

P-values

**Supplementary Figure 7: Resistance to IL1β driven demethylation in *Tet2*-KO progenitors promotes differential hypermethylation associated with cellular fate.**

WT and *Tet2*-KO mice were treated for 5 weeks with or without IL1β, purified BM derived LSK, CMP, and GMP cells were analyzed by whole genome bisulfite sequencing (n = 4 mice/group). **(a)** Experimental design, **(b)** differentially methylated regions (DMRs) from 50 CpGs tiling of the whole genome (>10% methylation difference, $q < 0.05$, as determined by SeqMonk) between vehicle or IL1β treatment in WT or *Tet2*-KO cells. **(c)** Top, MA plots showing methylation difference for LSK and CMP cells derived from *Tet2*-KO and WT mice treated with IL1β using literature-derived ChIP-seq binding sites of erythroid/megakaryocytic lineage specifying transcription factors (TFs). The percentage of sites with higher methylation in *Tet2*-KO than WT are shown in the top right corner of each graph. Bottom, the significance of motif enrichment using HOMER analysis tool within DMRs for TFs. With q values calculated by Homer. **(d)** As in Supplementary Fig. 7c, but showing lymphoid lineage specifying TFs. **(e)** Selected examples of genes showing methylation differences within promotor (top) or intragenic (bottom) regions. Student's two-tailed t-Test was used to determine significance except where otherwise specified in the figure legend: $^{*}p < 0.05$, $^{**}p < 0.01$, $^{***}p < 0.001$.

Supplementary Figure 8

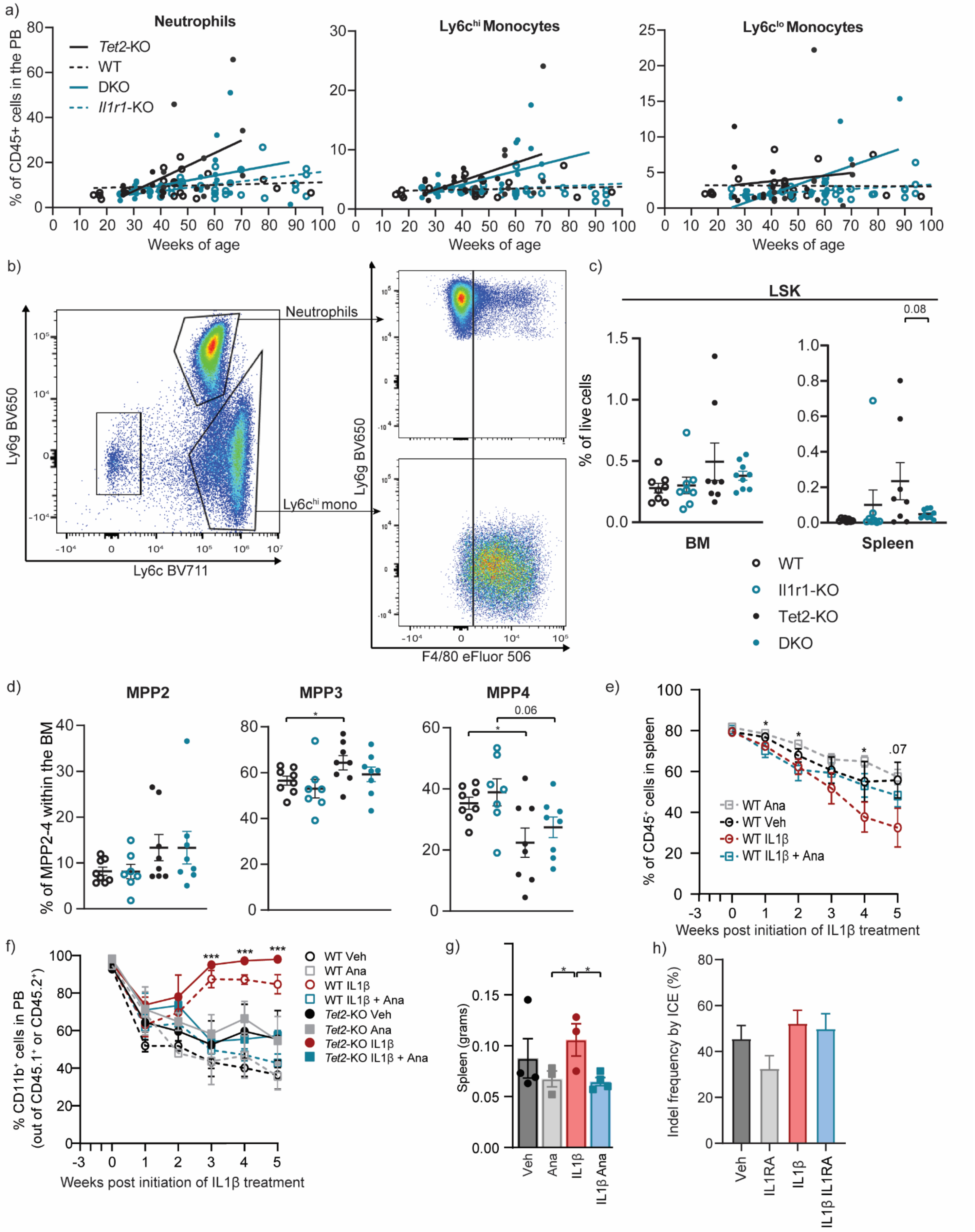

**Supplementary Figure 8: Disruption of IL1R1 signaling suppresses myeloid bias in *Tet2*-KO mice.**

**(a-d)** *Vav-cre Tet2$^{fl/fl}$* (*Tet2*-KO), *Il1r1$^{-/-}$* (*Il1r1*-KO), and *Vav-cre Tet2$^{fl/fl}$Il1r1$^{-/-}$* (double knockout; DKO) mice over 60 weeks of age. **(a)** Frequency of neutrophils, Ly6c$^{hi}$ monocytes, and Ly6c$^{lo}$ monocytes in the PB. **(b)** Representative gating within BM of the percentage of Ly6c$^{hi}$ monocytes/macrophages which are F4/80$^{+}$. **(c)** The frequency out of live cells of LSK cells within the BM and spleen. **(d)** Frequency out of live cells of MPP2, MPP3, and MPP4 out of MPP2-4 within the BM. **(e-g)** Lineage-depleted BM cells derived from WT CD45.1 and *Tet2*-KO CD45.2 mice were transplanted into lethally irradiated WT CD45.1/2 mice. 3 weeks later, mice were treated with IL1β (500 ng/mouse/day) or vehicle as well as with or without an IL1R1 antagonist (anakinra, 100mg/kg) (n = 3-4 mice/group). **(e)** Percentage of WT CD45.1 donor cells out of all CD45$^{+}$ cells (chimerism) in PB. **(f)** Percentage of WT CD45.1 and *Tet2*-KO CD45.2 donor cells which are myeloid in the PB. **(g)** Spleen weight in grams. **(h)** Indel frequency of individually picked colonies for vehicle, IL1RA, IL1β and IL1β with IL1RA is determined by Sanger sequencing for infer CRISPR edits (ICE) analysis. Error bars represent mean ± SEM. Student's two-tailed t-Test was used to determine significance: $^{*}p < 0.05$, $^{**}p < 0.01$, $^{***}p < 0.001$.
